# Working memory capacity is linked to directed exploration: behavioral, computational, and electrophysiological evidence

**DOI:** 10.64898/2026.11.08.765180

**Authors:** Dingding Duan, You Wang, Haowei Dai, Ruibin Zhang

## Abstract

Balancing the exploitation of known rewards against the exploration of uncertain options is central to adaptive decision-making, yet the cognitive limits on strategic exploration are poorly understood. Directed exploration, the deliberate sampling of uncertain options for information, may depend on working memory (WM), but whether WM capacity (WMC) is selectively related to it, and through which processes, is unknown. Fifty-two participants from the upper and lower tails of an operation-span distribution (26 high-WMC, 26 low-WMC) performed an adapted Horizon Task during EEG recording. We combined behavioral strategy classification, hierarchical Bayesian reinforcement learning, stimulus-locked event-related potentials (ERPs), and an exploratory Bayesian mediation analysis. High-WMC participants showed more directed exploration and less exploitation, with no group difference in random exploration. A model in which accumulated sampling history discounts option value best explained choices in both groups, and the discounting parameter ω was larger in the high-WMC group. Statistically, ω accounted for the WMC–directed exploration association (indirect effect *ab* = 0.64, 95% CrI [0.28, 1.05]). Frontocentral P2 amplitude (180–280 ms) during directed exploration was larger in the high-WMC group, and the P2 difference between directed exploration and exploitation correlated with behavioral preference for directed exploration (*r* = .29, *p* = .040). These findings suggest that individual differences in exploration may reflect not only the willingness to seek information but also the capacity to use what has been sampled. The WMC-related difference lay in information use rather than choice noise and was already visible in early option evaluation.

## Introduction

In uncertain environments, adaptive behavior requires organisms to balance two competing demands: exploiting options that provide reliable rewards and exploring alternatives that may yield greater future benefits (Sutton & Barto, 2018). This exploration–exploitation tradeoff represents a fundamental computational challenge for resource-limited systems, requiring decisions that integrate previously acquired value information with judgments about when exploring may provide useful new information. This problem is widespread across behavioral domains, including foraging, spatial exploration, learning, and memory-guided decision making (Hills et al., 2015), and is also central to artificial reinforcement-learning systems, in which agents must balance exploring alternative actions against exploiting actions with high estimated rewards (Auer et al., 2002).

In humans, exploration–exploitation strategies also vary substantially across individuals, and such variability has been linked to psychiatric symptoms (Addicott et al., 2017). In depression, symptom severity has been associated with differences in exploratory choice (Blanco et al., 2013). In schizophrenia, reduced directed exploration has been associated with anhedonia (Strauss et al., 2011), and patients have shown lower directed exploration than healthy controls without a comparable reduction in random exploration (Waltz et al., 2020), although findings vary across tasks and patient groups. Resolving the exploration–exploitation tradeoff requires integrating and maintaining multiple sources of information, such as the values and uncertainty of available options, under limited cognitive resources. These resources vary systematically across individuals (Conway et al., 2003). This raises the possibility that stable differences in cognitive resources may contribute to individual variability in exploration strategy, yet the mechanisms underlying this variability remain incompletely characterized.

Understanding why individuals differ in exploration strategies requires clarifying how exploratory choices use information. Although exploration may reduce immediate reward, it can provide information that improves future decisions under uncertainty (Cohen et al., 2007). In human behavior, this information can be acquired through different exploratory strategies. Individuals may increase the randomness of their choices to broaden the range of options they sample (random exploration), or selectively favor options expected to provide more informative outcomes (directed exploration). Although both strategies increase choices of alternatives beyond the currently preferred option, they may place different demands on cognitive resources (Wilson et al., 2014). Directed exploration in particular requires evaluating information associated with available options, including their expected value, uncertainty, and prior sampling history (Gershman, 2018). Using such information requires maintaining and integrating representations of previous experience to guide subsequent choices, making working memory (WM) a plausible cognitive mechanism underlying individual differences in directed exploration. WM provides a capacity-limited system for maintaining and manipulating task-relevant representations (Baddeley, 2012), and the reinforcement learning–working memory framework demonstrated that WM contributes independently of incremental reinforcement learning processes and that its contribution is constrained by capacity limitations and task demands (Collins & Frank, 2012; Collins, 2025).

Extending this framework to exploratory decision-making, recent work showed that experimentally increasing memory demands specifically reduced directed exploration in a multi-armed bandit task (Brown et al., 2022). These findings suggest that WM limitations may alter exploration strategy selection by constraining the maintenance and integration of option-specific information during uncertain decisions, but they leave open whether stable individual differences in working memory capacity (WMC) are associated with exploration strategy and, if so, through which computational process.

Existing causal evidence linking WM to exploration has relied primarily on transient manipulations of memory demands within a task session (Brown et al., 2022; Cogliati Dezza et al., 2019). WMC has traditionally been conceptualized as a domain-general individual-difference construct reflecting controlled attention and the ability to maintain task-relevant representations under interference (Engle, 2002). Recent psychometric work found that latent measures of directed and value-guided exploration were associated with WMC (Witte et al., 2025), providing initial evidence for an association between WMC and exploration. The same study, however, reported poor to moderate reliability and minimal cross-task correlations for exploration measures, a limitation of task-based individual-difference measures more generally (Zorowitz & Niv, 2023), and questioned whether the exploration factors provide information beyond general performance-related constructs such as WMC. No computational process linking the two constructs has yet been identified. One candidate process is the use of prior sampling history. If higher WMC facilitates the use of sampling history in value computation, individuals may more strongly discount extensively sampled options, increasing choices of less-sampled alternatives. Because the cognitive computations underlying exploration cannot be directly observed from choice behavior alone, computational modeling provides a framework for identifying latent processes underlying individual differences in exploratory strategies. Within this framework, information discounting characterizes exploration as a process in which options lose informational value as experience accumulates (Cogliati Dezza et al., 2017), allowing the influence of prior sampling history on option valuation to be quantified without requiring an explicit uncertainty term. We therefore asked whether WMC is associated with exploration not only through overall strategy preference but also through how prior sampling information is incorporated into value computation.

A related question concerns when WMC-related differences in exploration emerge during the decision process. Neuroimaging and brain stimulation studies have begun to identify the neural systems supporting uncertainty-guided exploration, implicating the rostrolateral and frontopolar prefrontal cortex (Badre et al., 2012; Zajkowski et al., 2017) and fronto-striatal circuits (Cockburn et al., 2022; Wyatt et al., 2024). However, these approaches provide limited temporal resolution and cannot reveal the temporal dynamics of exploratory computations. Electroencephalography (EEG), through event-related potential (ERP) analysis, offers millisecond-level temporal resolution to address this question. Existing EEG studies of exploration have identified neural correlates at different stages of the decision process, with particular attention to responses following choice and outcome presentation. Exploratory decisions have been associated with enhanced feedback-locked P300 responses (Hassall et al., 2013, 2019), while response-locked medial frontal theta activity has been linked to uncertainty during exploratory choices (Cavanagh et al., 2012). These findings leave open when exploration-relevant information is first evaluated during the unfolding decision. This question is particularly relevant for directed exploration, which requires integrating expected value with potential information gain at the moment of choice. The frontocentral P2, an early stimulus-locked ERP component associated with task-relevant stimulus evaluation (Potts, 2004; Potts & Tucker, 2001), provides a candidate marker of early option evaluation. A related prefrontal P2 component has been linked to uncertainty about stimulus–response mappings (Aydin et al., 2024), making P2 relevant to directed exploration, which requires evaluating the informational relevance of available options before choice. Top-down modulation of early visual responses during WM encoding is related to subsequent WM performance (Zanto & Gazzaley, 2009), and increased WM load reduces P1 amplitude (Pratt et al., 2011). These findings suggest that WM-related processes can be reflected in early stimulus processing, but whether individual differences in WMC are detectable in early option evaluation during directed exploration remains unknown.

To address these gaps, the present study integrates behavioral, computational, and electrophysiological approaches to examine whether individual differences in WMC are associated with exploratory decision making and, if so, at which processing stage. We focused on information discounting as a computational measure of how prior sampling history influences option valuation, and on stimulus-locked P2 activity as an early marker of option evaluation. We hypothesized that higher WMC would be associated with a greater proportion of directed exploration (Hypothesis 1) and with stronger information discounting of previously sampled options (Hypothesis 2), and that WMC-related differences in directed exploration would be detectable in P2 amplitude during early option evaluation (Hypothesis 3). As a secondary question, we tested whether information discounting statistically accounted for the association between WMC and directed exploration. Together, these analyses characterize WMC-related differences in exploration at behavioral, computational, and neural levels.

## Materials and Methods

### Participants and Design

The study employed a between-subject extreme-groups design based on individual differences in WMC, combined with a within-subject decision-making paradigm. WMC was assessed in a screening sample (*N* = 200) with the Automated Operation Span Task (Mei et al., 2021).

Participants were selected from the upper (≥85th percentile) and lower (≤15th percentile) tails of the distribution to maximize sensitivity to individual differences in cognitive capacity. Of the 30 individuals eligible in each tail, 26 per group agreed to participate and completed the full session, yielding a final sample of 52 healthy young adults aged 19–23 years (*M* = 21.24). The High-WMC group included 5 male and 21 female participants, and the Low-WMC group included 7 male and 19 female participants. All participants had normal or corrected-to-normal vision and no history of neurological or psychiatric disorders. No participants were excluded from the analyses.

### Sample size justification

The sample size was set by the number of eligible screened individuals in each tail of the WMC distribution who agreed to participate. A sensitivity analysis for independent-samples *t* tests (26 per group, α = .05, two-tailed, power = .80) showed that the design could detect group differences of Cohen’s *d* ≥ 0.79; smaller effects may have gone undetected.

### Ethics

The study was approved by the Biomedical Ethics Committee of Southern Medical University (approval no. Nan Yi Lun Li Shen [2026] No. 35) and conducted in accordance with the Declaration of Helsinki. All participants provided informed consent.

### Decision-Making Task and Behavioral Measures

The decision-making task (Figure 1) was based on the Horizon Task (Wilson et al., 2014), as adapted by Cogliati Dezza et al. (2017). The original design was proportionally reduced from 128 to 90 blocks to shorten the session while maintaining condition balance. Each block comprised a six-trial forced-choice phase, in which participants selected among three card decks according to program-determined instructions, followed by a free-choice phase (one to six trials, exponentially distributed and never cued) in which they chose freely among the same decks to maximize cumulative reward. Three new decks were used on every block.

**Figure 1.**
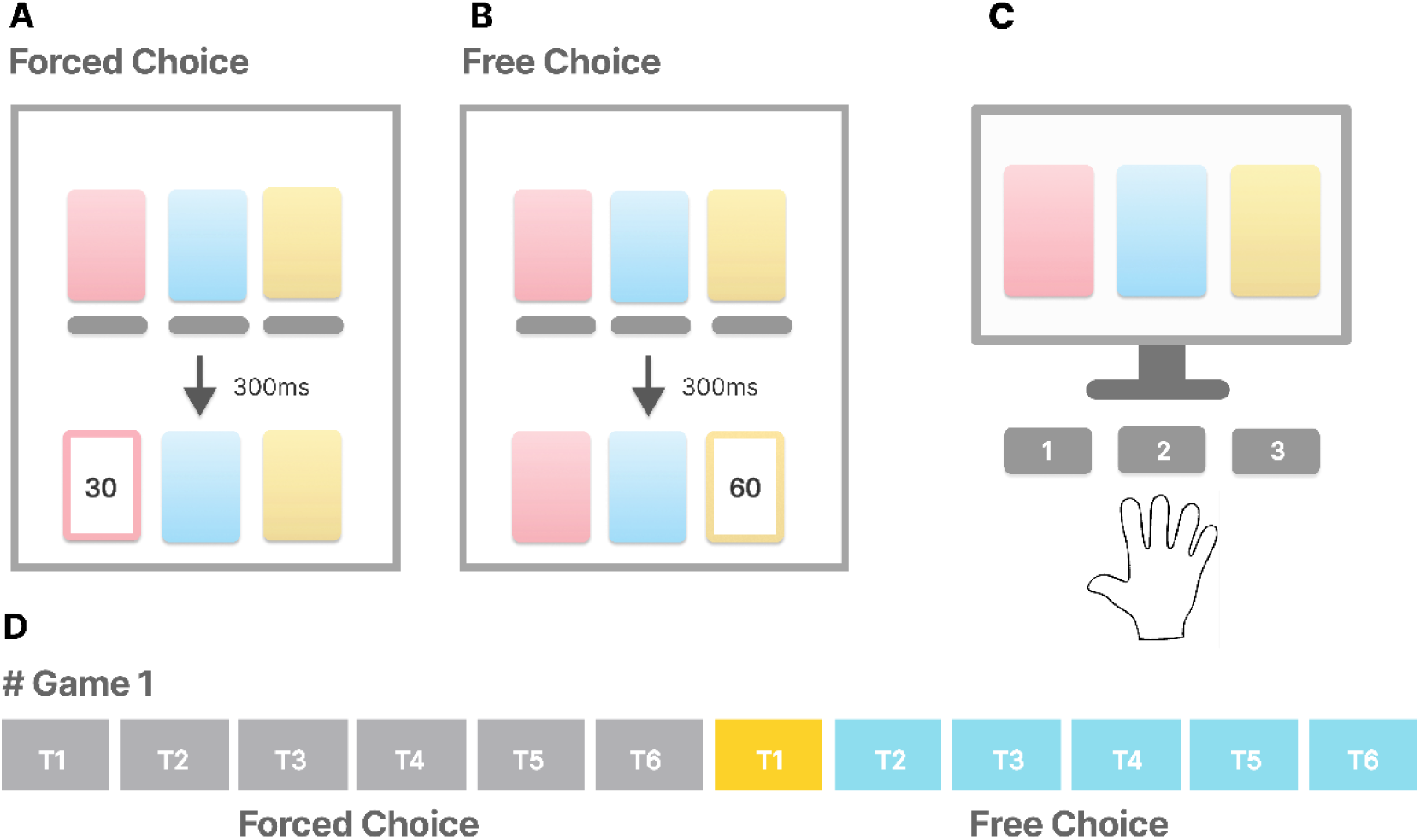
Behavioral paradigm. The task comprised a forced-choice phase followed by a free-choice phase. (A) Forced-choice phase: participants selected among three card decks (red, blue, and yellow) according to predefined instructions and received reward feedback after each choice; sampling structure differed between the equal- and unequal-information conditions. (B) Free-choice phase: participants made autonomous choices based on acquired information. (C) Response mapping: three keyboard keys were used to select among decks. (D) Task structure: each block comprised a forced-choice phase followed by a free-choice phase of variable length. Behavioral analyses focused on the first free-choice trial (highlighted in yellow).

Two factors were manipulated orthogonally during the forced-choice phase. Information was varied through the sampling schedule, with each deck sampled either twice under equal information or four, two, and zero times under unequal information. Reward was varied independently. On each block, reward magnitude was drawn from a truncated Gaussian distribution centered on a generative mean of 30 points under a low-reward context or 50 points under a high-reward context. When deck rewards were unequal, individual deck means were offset from this block-level mean by 0, 4, 12, or 20 points, producing a 2:1 split in which two decks received lower values and one deck a higher value, or one deck a lower value and two decks higher values, with the specific deck assigned the odd-one-out value randomized on each block.

Crossing the information and reward manipulations yielded four block types totaling 90 blocks, with block order randomized for each participant. The Information condition combined unequal information with equal reward and comprised 30 blocks. The Reward condition combined equal information with unequal reward and likewise comprised 30 blocks. The Baseline condition combined equal information with equal reward and comprised 14 blocks. The Mixed condition combined unequal information with unequal reward and comprised 16 blocks. The Information and Reward conditions were the primary conditions of theoretical interest, while the Baseline and Mixed conditions served as controls. Within each condition, blocks were divided evenly across the low-and high-reward contexts or, where applicable, across the two possible 2:1 assignment.

Stimuli were presented with PsychoPy, with participants seated 60 cm from the monitor. A trigger marking the onset of the card array was sent to the EEG system on each trial.

### Behavioral Data Processing and Analysis

Behavioral data were processed using custom MATLAB scripts. Analyses focused on the first free-choice trial in each block to isolate initial decision policies prior to within-block learning effects. Trials in which participants did not follow forced-choice instructions were excluded to preserve the intended information structure. Each first free-choice decision was classified into three mutually exclusive strategies based on sampling and reward history: directed exploration (selecting an unsampled option), exploitation (selecting the option with the highest observed mean reward), and random exploration (all remaining choices). For each participant, strategy proportions were computed and entered into a 2 (WMC group: High-WMC, Low-WMC) × 3 (strategy: directed exploration, exploitation, random exploration) mixed-design ANOVA using the aligned rank transform procedure (Wobbrock et al., 2011) in the *ARTool* package. Follow-up independent-samples *t* tests were Bonferroni-corrected. Effect sizes are reported as partial eta squared (ηp^2^) for ANOVAs and Cohen’s *d* for *t* tests.

### Computational Modeling

To identify the computational processes underlying exploration–exploitation behavior, we implemented three progressively extended reinforcement learning models adapted from Cogliati Dezza et al. (2017): a standard reinforcement learning model (sRL), a knowledge reinforcement learning model (kRL), and a gamma-knowledge reinforcement learning model (gkRL). The models extend reward-based learning by incorporating accumulated information about each option and allowing its influence on decision value to vary nonlinearly.

The sRL model updates the expected value of each option according to the Rescorla–Wagner learning rule:

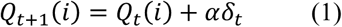

where *α* is the learning rate and *δ_t_* = *R_t_* − *Q_t_*(*i*) is the reward prediction error. Choice probabilities were generated using a softmax function:

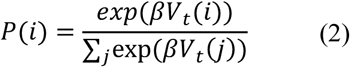

where *β* is the inverse-temperature parameter controlling choice stochasticity. For sRL, the decision value *V_t_*(*i*) corresponds to the learned reward value *Q_t_*(*i*).

The kRL model extends sRL by additionally tracking information accumulated through previous selections of each option. Specifically, accumulated information *I_t_*(*i*) was defined as the cumulative number of times option *i* had been selected up to trial *t*, such that 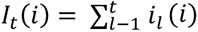, here *i_l_*(*i*) if option *i* as selected on trial *l*, and 0 otherwise. This information term as incorporated into decision value as an information-based suppression term eighted by *ω*. Building on kRL, the gkRL model allows accumulated information to influence decision value nonlinearly:

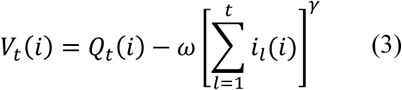

where *ω* controls the strength of information-based suppression and *γ* determines the nonlinear mapping of accumulated information onto decision value. When *γ* = 1, the gkRL formulation reduces to the linear information term used in kRL; *γ* < 1 and *γ* > 1 indicate sublinear and superlinear effects of accumulated information on decision value, respectively.

Models were fitted separately to the High-WMC and Low-WMC groups using hierarchical Bayesian estimation, with participant-level parameters estimated within group-level distributions. The three model variants were compared within each group (six fitted models in total) using the Widely Applicable Information Criterion (WAIC; Watanabe, 2010), with lower values indicating better expected out-of-sample predictive performance. Parameter recovery and posterior predictive checks (PPCs) were conducted for the best-fitting model to assess parameter identifiability and model validity.

### EEG Data Acquisition and Event-Related Potential Analysis

EEG data were recorded with a 64-channel BrainRecorder system arranged according to the international 10–20 system, digitized at 500 Hz, with FCz as the online reference and AFz as ground. Electrode impedances were kept below 5 kΩ throughout recording. Offline preprocessing was conducted in MATLAB (R2019b) using EEGLAB (version 14.1.2) (Delorme & Makeig, 2004) and ERPLAB (Lopez-Calderon & Luck, 2014). Data were bandpass filtered (0.1–30 Hz) and re-referenced offline to the average of TP9 and TP10. Artifact correction was performed using ICA, with components identified via ICLabel (Pion-Tonachini et al., 2019) and ADJUST (Mognon et al., 2011) and removed when classified as artifactual with high confidence, followed by visual inspection. Bad channels were interpolated using spherical spline interpolation. Following artifact rejection, an average of 6.04% of trials were excluded per participant.

ERP analyses focused on the P2 component, a frontocentral positivity associated with early evaluation of task-relevant stimuli. Epochs (−200 to 800 ms) were time-locked to the onset of the card array on the first free-choice trial of each block and baseline-corrected using the −200 to 0 ms pre-stimulus interval. Mean amplitude within 180–280 ms was used as the dependent measure. Analyses targeted a frontocentral region of interest consistent with prior literature and the observed scalp distribution in the present dataset. FCz was used as the representative electrode for univariate analyses. A 2 × 3 mixed-design ANOVA was conducted on P2 amplitudes, with working memory group as the between-subjects factor and decision strategy as the within-subjects factor. Effect sizes are reported as generalized eta squared (ηG^2^). Feedback-locked ERPs were also analyzed (Supplementary Materials 2).

To further examine whether these P2 amplitude differences corresponded to individual differences in decision behavior, a behavioral index was defined as the difference in selection proportion between directed exploration and exploitation, and a corresponding neural index was defined as the difference in P2 amplitude between the same two conditions at electrode FCz. The association between these two indices was assessed using Pearson correlation.

### Mediation Analysis of WMC, Information Discounting, and Exploration Strategy

To test whether WMC was statistically associated with exploration strategy through the computational parameter ω, we conducted an exploratory Bayesian hierarchical mediation analysis of the pathway WMC group → ω → strategy. WMC was coded as a binary group variable, ω was natural-log transformed, and strategy (see Behavioral Data Processing and Analysis) was the three-level categorical outcome. Strategy was classified from task-defined variables independent of model output, whereas ω was estimated from the gkRL model, so the two measures did not share a statistical source despite deriving from the same task. Trials coded as invalid choices (*n* = 43) were excluded from 52 participants. Because strategy was measured repeatedly within participants, all models predicting strategy included a participant-level random intercept.

Four models were estimated separately with *brms*. Path A modeled log(ω) as a function of WMC group (Gaussian regression). Path B modeled strategy as a function of log(ω) (categorical mixed-effects regression; *b* denotes the directed-exploration contrast and *β* the random-exploration contrast, both relative to exploitation). Path C estimated the total effect of WMC group on strategy, and Path C′ the direct effect of WMC group controlling for log(ω). The indirect effect was the product of the posterior draws of *a* and *b* at each iteration, which assumes independence of the two posteriors, plausible because the models share no parameters and operate at different levels of analysis. It is expressed on the log-odds scale and summarized by its posterior mean, 95% credible interval (CrI), and the posterior probability of exceeding zero. All models used 4 chains of 4,000 iterations (2,000 warm-up), yielding 8,000 post-warm-up draws; convergence was confirmed (R̂ = 1.00).

## Results

### Working Memory Capacity and Exploration–Exploitation Strategy Preferences

We first examined the decision strategy employed on the first free-choice trial under the unequal-information condition (Figure 2). Strategy distributions differed markedly between the High-WMC and Low-WMC groups (Figure 2A). The aligned-rank-transform ANOVA revealed a significant strategy × group interaction, *F*(2, 100) = 10.14, *p* < .001, ηp^2^ = .17, whereas the main effect of group was not significant (*p* = .98). Bonferroni-corrected *t* tests showed that the Low-WMC group exhibited a higher proportion of exploitation than the High-WMC group, *t*(50) = 2.65, *p* = .032, *d* = 0.73, whereas the High-WMC group showed a higher proportion of directed exploration, *t*(50) = −2.63, *p* = .034, *d* = 0.73. The groups did not differ in random exploration (*p* = .74). A count-based quasibinomial model accounting for the number of valid trials yielded a consistent pattern (exploitation: *p* = .012; directed exploration: *p* = .013; random exploration: *p* = .27).

**Figure 2.**
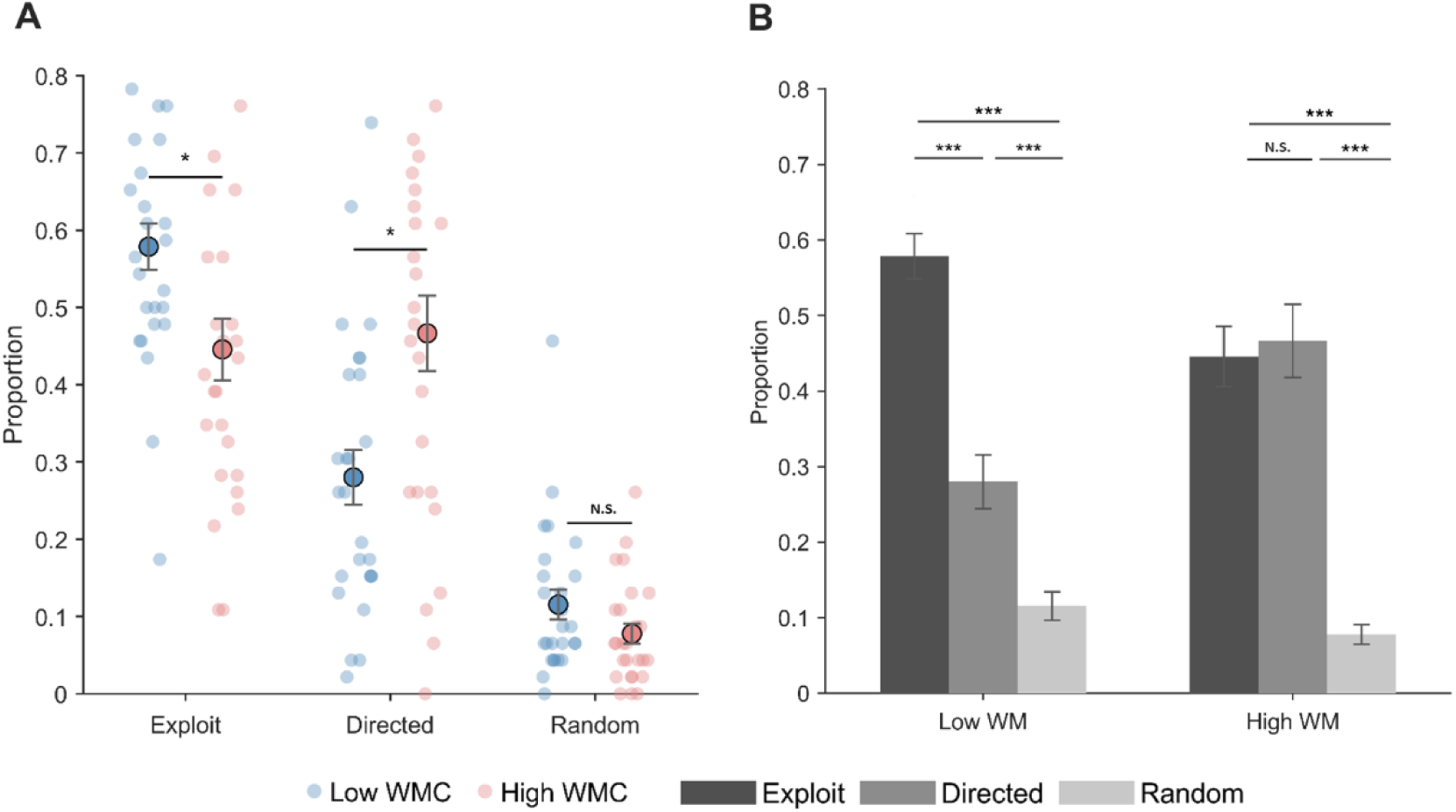
Strategy preferences on the first free-choice trial of unequal-information blocks. (A) Individual strategy proportions by WMC group, with small dots representing individual participants and large circles indicating group means. The High-WMC group showed greater directed exploration and less exploitation than the Low-WMC group, with no significant group difference in random exploration. (B) Strategy profiles within each WMC group, with brackets indicating within-group pairwise comparisons. Error bars represent ±1 *SEM*. Group differences were assessed using an aligned rank transform mixed ANOVA followed by Bonferroni-corrected post hoc tests. * *p* < .05, *** *p* < .001; N.S., not significant.

Strategy proportions differed within both groups (Figure 2B): High-WMC, *F*(2, 50) = 35.42, *p* < .001, ηp^2^ = .59; Low-WMC, *F*(2, 50) = 56.58, *p* < .001, ηp^2^ = .69. The Low-WMC group used exploitation more often than directed exploration, and directed exploration more often than random exploration (all adjusted *p*s < .001). The High-WMC group showed no difference between exploitation and directed exploration (*p* = 1.00), although both were used more often than random exploration (both adjusted *p*s < .001). The group difference in directed exploration held in both reward contexts (low reward, High vs. Low WMC: .582 vs. .428, *p* = .040; high reward: .331 vs. .146, *p* = .005), and the Low-WMC group showed more exploitation in both. Both groups adjusted similarly to reward variation, with directed exploration decreasing and exploitation increasing in the high-reward context.

### Computational Modeling Results

The three models (sRL, kRL, gkRL) were compared by WAIC within each group (Table 1). Chains converged for all models (maximum R̂ = 1.01, adequate effective sample sizes, no divergent transitions). The gkRL model fit best in both groups (High-WMC: WAIC = 10,789.7 vs. 11,557.5 for kRL, ΔWAIC = 767.8; Low-WMC: 11,813.3 vs. 11,971.1, ΔWAIC = 157.8), with the rank order gkRL > kRL > sRL in both. Because models were fitted separately by group, WAIC values are comparable only within a group. Posterior predictive checks showed that gkRL reproduced the observed proportions of exploitation, directed exploration, and random exploration in both groups (*r*s = .56–.79, *p*s < .01; RMSEs = 0.056–0.173), so gkRL was used for all subsequent analyses.

**Table 1.** WAIC Comparison of Six Candidate Models (Fitted Separately Within Each WMC Group)

| **Model** | **elpd_waic (SE)** | **WAIC** | **p_waic** |
|---|---|---|---|
| gkRL_High | −5394.8 (84.4) | 10789.7 | 79.2 |
| kRL_High | −5778.8 (87.0) | 11557.5 | 71.8 |
| sRL_High | −5848.8 (87.8) | 11697.6 | 52.8 |
| gkRL_Low | −5906.6 (86.4) | 11813.3 | 68.4 |
| kRL_Low | −5985.5 (86.8) | 11971.1 | 60.5 |
| sRL_Low | −5995.5 (87.0) | 11991.0 | 49.9 |

**Note.** elpd\_waic denotes the expected log pointwise predictive density; larger values indicate better fit. WAIC is its negative double, with smaller values indicating superior fit. p\_waic denotes the effective number of parameters. High and Low refer to the high working memory capacity (High-WMC) and low working memory capacity (Low-WMC) groups, respectively. WAIC values are comparable only among models fitted to the same group.

Parameter estimates from the gkRL model are shown in Figure 3. The learning rate (α) and inverse temperature (β) did not differ between groups (both *p*s > .05). In contrast, the information suppression weight ω was higher in the High-WMC group (*M* = 8.22) than in the Low-WMC group (*M* = 2.51), *t*(32.28) = 2.90, *p* = .007, *d* = 0.80, indicating that higher-WMC participants more strongly discounted the value of previously sampled options. The nonlinear exponent γ also differed between groups, *t*(29.45) = −10.12, *p* < .001, but its parameter recovery was poor in both groups (High-WMC: *r* = .22; Low-WMC: *r* = −.18), so this difference was not interpreted.

**Figure 3.**
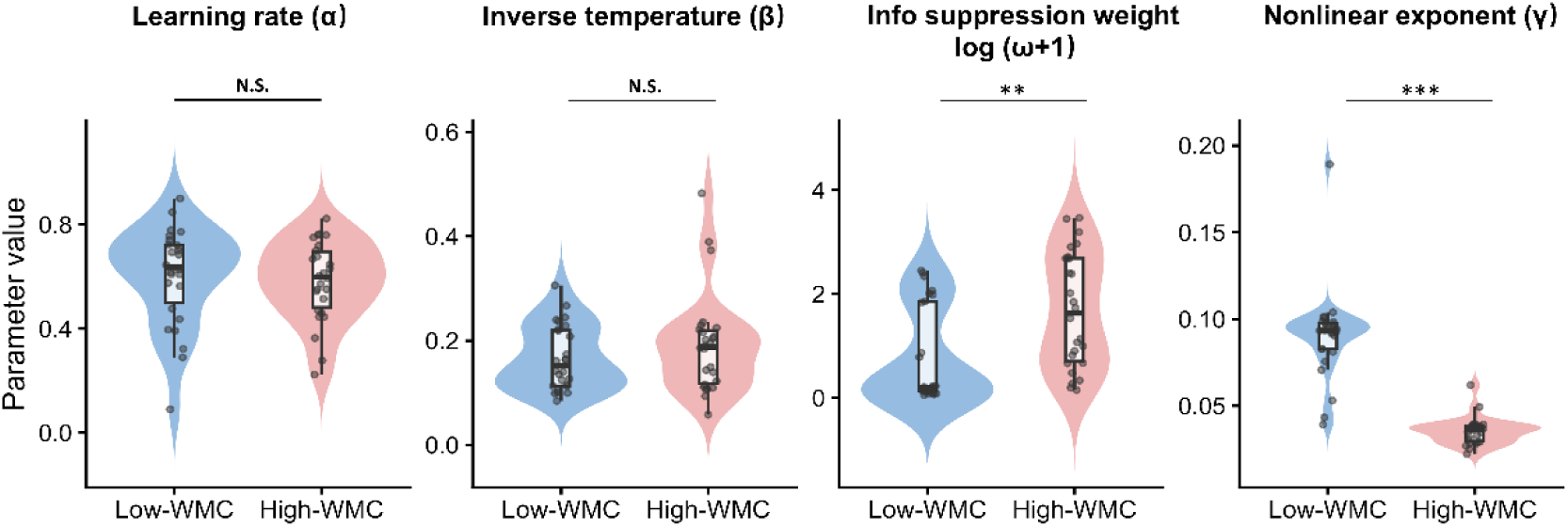
Distributions of gkRL parameters by WMC group. Violin plots show individual-level posterior estimates of the learning rate (α), inverse temperature (β), information suppression weight (ω), and nonlinear exponent (γ). The values of ω are plotted on a log(ω + 1) scale to account for its right-skewed distribution. Boxplots indicate the median and interquartile range, and points represent individual participants. Statistical significance is indicated by ** *p* < .01 and *** *p* < .001, whereas N.S. denotes a nonsignificant difference. The parameter γ showed poor recovery in both groups and should therefore be interpreted with caution.

### Working Memory Capacity and P2 Amplitude Depend on Decision Strategy

Neural analyses focused on the first free-choice trial and examined the P2 component within the 180 to 280 ms post-stimulus window (Figure 4A). There was a main effect of group, *F*(1, 50) = 8.19, *p* = .006, ηG^2^ = .066, with larger overall P2 amplitude in High-WMC (*M* = 3.66 μV, *SEM* = 0.46) than Low-WMC participants (*M* = 1.80 μV, *SEM* = 0.46). Critically, the group × strategy interaction was significant, *F*(2, 100) = 5.65, *p* = .005, ηG^2^ = .061, indicating that WMC-related differences in early neural processing varied across strategies. Across groups, P2 was largest for directed exploration (*M* = 3.77 μV, *SEM* = 0.62) and significantly exceeded P2 for random exploration (*M* = 2.04 μV, *SEM* = 0.57, *p* = .027). Directed exploration did not differ significantly from exploitation (*M* = 2.38 μV, *SEM* = 0.35, *p* = .11), nor did exploitation from random exploration (*p* = 1.00).

**Figure 4.**
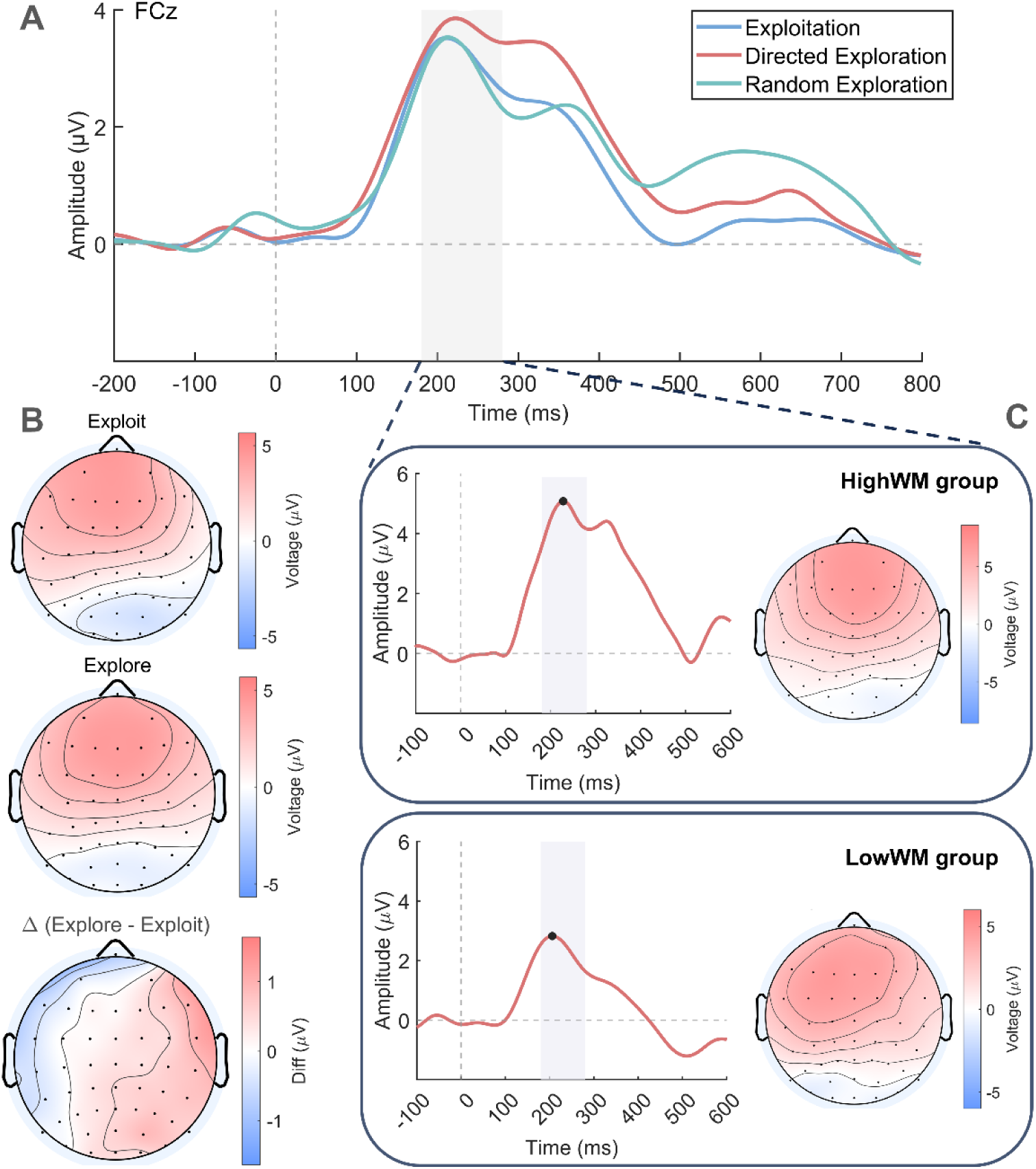
P2 results at FCz by decision strategy and WMC group. (A) Grand-average ERPs for exploitation, directed exploration, and random exploration across all participants; 0 ms is card-array onset and the gray band marks the 180–280 ms analysis window. (B) Scalp topographies (180–280 ms) for exploitation and directed exploration, and their difference (directed exploration − exploitation). (C) Directed-exploration ERP waveforms at FCz and scalp topographies for the High-WMC (top) and Low-WMC (bottom) groups; black dots mark the P2 peak. Waveforms were smoothed for display only; statistics used unsmoothed data.

Scalp topographies for 180–280 ms (Figure 4B) showed a frontocentral positivity for both exploitation and directed exploration. The difference topography (directed exploration minus exploitation) was small (maximum ≈ 1.5 μV) and spatially diffuse, with its largest positive values over right frontal sites

The interaction was driven by directed exploration (Figure 4C). During directed exploration, P2 amplitude was larger in High-WMC (*M* = 5.84 μV, *SEM* = 0.75) than in Low-WMC participants (*M* = 1.90 μV, *SEM* = 0.81; estimate = −4.14, *t*(144) = −4.18, *p* < .001). Group differences were not significant for exploitation (*p* = .84) or random exploration (*p* = .100).

### Relationship Between P2 Amplitude and Individual Strategy Preferences

To examine whether the P2 amplitude differences reported above were related to individual differences in decision behavior, the association between behavioral strategy preference and neural activity was analyzed. A behavioral index was constructed as the difference in selection proportion between directed exploration and exploitation. A corresponding neural index was defined as the difference in P2 amplitude between the directed exploration and exploitation conditions at electrode FCz. Pearson correlation analysis revealed a significant positive association between these two indices, *r*(50) = .29, 95% CI [.02, .52], *p* = .040 (Figure 5A).

**Figure 5.**
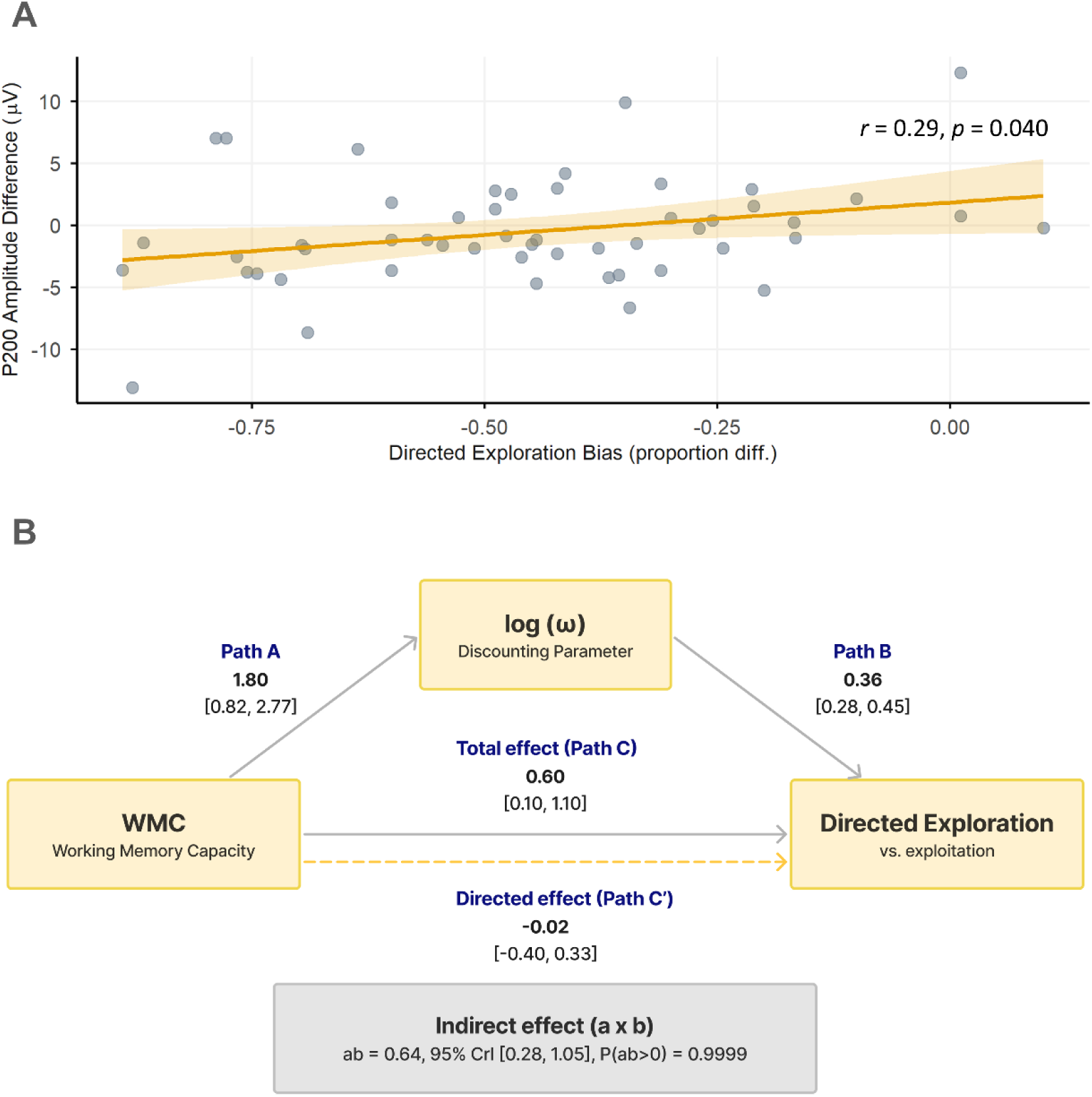
Early neural correlates of strategy preference and Bayesian mediation by information discounting. (A) Association between the behavioral index, calculated as the difference in directed exploration and exploitation selection proportions, and the neural index, defined as the difference in P2 amplitude between these conditions at FCz. Each point represents one participant. The line indicates the linear fit with its 95% confidence interval (*r* = .29, *p* = .040). (B) Bayesian mediation analysis of the association between WMC group and directed exploration relative to exploitation through log(ω). Solid gray arrows indicate posterior mean coefficients with 95% credible intervals (CrIs), and the dashed arrow represents the direct effect after controlling for log(ω) (Path C′). Path A was estimated using Gaussian regression, whereas Path B and the total effect (Path C) were estimated using categorical mixed-effects regression with exploitation as the reference category and participant-level random intercepts. The gray box presents the indirect effect (*a* × *b*), its 95% CrI, and the posterior probability that the indirect effect is positive, *P*(*ab* > 0).

### Information Discounting Statistically Accounts for the WMC and Directed Exploration Association

To examine whether the information-discounting parameter *ω* statistically accounted for the association between working memory capacity and exploration strategy selection, we conducted an exploratory Bayesian mediation analysis (Figure 5B). The High-WMC group showed higher log(ω) than the Low-WMC group (Path A: *a* = 1.80, 95% CrI [0.82, 2.77]), and log(ω) was associated with greater odds of directed exploration relative to exploitation (Path B: *b* = 0.36, 95% CrI [0.28, 0.45]; *OR* = 1.43, 95% CrI [1.32, 1.56]; Figure 5B). log(ω) was not clearly associated with random exploration relative to exploitation (*β* = −0.07, 95% CrI [−0.16, 0.03]).

The total effect of WMC group on directed exploration relative to exploitation was positive (Path C: *c* = 0.60, 95% CrI [0.10, 1.10]). When log(ω) was added to the model, the direct effect was attenuated to near zero (Path C′: *c*′ = −0.02, 95% CrI [−0.40, 0.33]), while log(ω) remained positively associated with directed exploration (*b* = 0.36, 95% CrI [0.27, 0.46]).

The indirect effect of WMC group on directed exploration through log(ω) was positive (*ab* = 0.64, 95% CrI [0.28, 1.05]; posterior probability that *ab* > 0 = .9999). The total effect was thus substantially reduced once log(ω) was accounted for. This pattern is consistent with an indirect statistical association between WMC and directed exploration through ω, not with evidence of causal mediation.

## Discussion

We examined how working memory capacity (WMC) is associated with exploration strategy using behavioral, computational, electrophysiological, and mediation analyses. Higher-WMC individuals showed more directed exploration and stronger information discounting of previously sampled options (larger ω), and exploratory mediation analysis indicated that ω statistically accounted for the association between WMC and directed exploration. A stimulus-locked frontocentral P2 was larger during directed than random exploration, and P2 amplitude during directed exploration was larger in the high-WMC group. These findings suggest that WMC-related differences in directed exploration are associated with how sampled information enters option valuation and are already detectable during early option evaluation.

The behavioral pattern suggests that the greater exploratory choice in the high-WMC group reflected directed rather than random exploration. Group differences emerged for directed exploration and exploitation, but not random exploration, while the choice-consistency parameter β did not differ between groups. Thus, the group difference was accompanied by differences in information discounting, but not in general choice consistency. The lack of a WMC effect on random exploration fits evidence that random exploration relies on partly separate mechanisms, including inhibitory activity in right dorsolateral prefrontal cortex (Chizari et al., 2025; Toghi et al., 2024). Earlier work shows that increasing cognitive demands changes how adults explore, but the direction differs across studies. In a multi-armed bandit task, higher memory demands reduced directed exploration (Brown et al., 2022). In dual-task studies, working memory load increased broad information sampling and shifted adults’ attention toward patterns seen in children (Sloutsky et al., 2026; Wan & Sloutsky, 2026). Differences in task structure and in how exploration is measured may explain this. Our findings concern stable capacity and not load. They suggest that individual differences in cognitive capacity are also associated with stable differences in the use of information during exploratory choice.

The computational results further specify the relationship between WMC and exploration strategy. The winning gkRL model represents directed exploration through discounting the value of previously sampled options as a function of sampling history, with ω indexing the strength of this discounting (Cogliati Dezza et al., 2017). Computationally cheaper heuristics have been proposed to complement more demanding uncertainty-guided exploration strategies (Dubois et al., 2021). This distinction is important for interpreting the role of WMC. Rather than indicating that higher-WMC individuals compute uncertainty more precisely, the present association between WMC and ω suggests that cognitive capacity may be related to how information accumulated across previous sampling is incorporated into subsequent choices. This interpretation is consistent with computational accounts in which selectively maintaining value information helps resolve the exploration–exploitation tradeoff (Hallquist & Dombrovski, 2019) and with resource-rational accounts in which cognitive constraints shape the strategies used to make decisions (Lieder & Griffiths, 2020). People also rely more on their past choices under cognitive load (Lai & Gershman, 2024), which may reduce the sampling of less-visited options. Recurrent networks with larger capacity likewise show more directed exploration (Flimm et al., 2026), although they also show less random exploration, which we did not observe in humans. The indirect effect was positive, while the residual direct association between WMC and directed exploration was centered near zero after accounting for ω, a pattern consistent with full statistical mediation. This suggests that individual differences in WMC may be reflected in directed exploration through differences in how previously sampled information is incorporated into subsequent choices. Because WMC was not experimentally manipulated, however, this statistical mediation does not by itself establish a causal pathway from WMC to ω to exploration, nor does it establish that ω specifically reflects WM maintenance.

Although gkRL outperformed kRL in model comparison, the nonlinear exponent γ was poorly identified. Posterior means of γ were close to zero in both groups (see Supplementary Materials 3), and recovery was poor. Recovery was good to excellent for α, β, and ω. The group difference in γ was therefore not interpreted. This limits conclusions about the functional form of information accumulation, but not the evidence for information discounting captured by ω. More variation in sampling frequency may be needed to estimate γ reliably. Task-derived measures also often show limited reliability (Zorowitz & Niv, 2023), so the ω findings should be replicated in independent samples.

The neural findings provide complementary evidence regarding when strategy-related processing occurs. P2 amplitude was larger during directed exploration than during random exploration (the difference from exploitation was not significant), and P2 amplitude during directed exploration was larger in the high-WMC group. The frontocentral P2 has been associated with early evaluation of task-relevant stimuli (Potts, 2004; Potts & Tucker, 2001), providing a plausible temporal marker for early option evaluation. The present findings extend this literature by showing that P2 activity differs across exploration strategies and that WMC-related differences were specific to directed exploration. These converging results suggest that strategy-relevant differences are detectable during the initial evaluation of choice options, before feedback is presented.

A supplementary analysis of feedback-locked activity (see Supplementary Materials 2) provides a further temporal distinction. The P300 elicited after outcome presentation also differentiated exploration strategies, consistent with previous reports of feedback-related neural correlates of exploration (Hassall et al., 2019), but this strategy difference was not modulated by WMC. The combination of a WMC-sensitive P2 during option presentation and a WMC-insensitive P300 during feedback is therefore consistent with a more pronounced role of WMC during early evaluation of choice-relevant information than during subsequent outcome processing. This pattern extends the reinforcement learning–working memory framework, which has emphasized the contribution of WM to maintaining information that supports learning alongside incremental reinforcement learning (Collins & Frank, 2012; Collins, 2025), by suggesting that capacity-related differences may also be expressed before feedback, when information is evaluated for the current choice. However, the absence of a WMC × P300 interaction does not establish that WMC has no influence on outcome processing.

Across levels, the findings provide converging but not identical evidence. The mediation analysis showed that ω statistically accounted for the association between WMC and directed exploration, identifying information discounting as a computational correlate of the WMC-related difference in exploration. In parallel, the P2 findings provide temporal evidence that strategy-related processing occurs during early option evaluation. These findings therefore converge on the involvement of information processing before choice, but they do not establish that P2 activity directly reflects ω. Because ω and P2 were not incorporated into a joint model, future work should test this computational-neural relationship more directly by integrating neural measures into computational models of exploration. If the capacity-related differences in ω generalize across tasks and samples, ω could serve as a marker of how stable capacity relates to exploration. This remains to be tested.

Several limitations qualify these conclusions. The present study intentionally selected participants from the upper and lower ends of the WMC distribution to increase sensitivity to stable individual differences. Accordingly, the present findings most directly support differences between higher-and lower-WMC groups and do not establish whether these effects vary linearly across the full WMC continuum (Preacher et al., 2005). The mediation analysis was based on separately estimated path models and therefore provides evidence for statistical mediation rather than establishing a causal mediation pathway; the indirect effect was also specific to the directed-exploration contrast and expressed on the log-odds scale. A supplementary check (see Supplementary Material 6) indicated that rescaling effects from the nested logistic models made only a minor contribution to the observed attenuation. Future work could further optimize the computational model to more fully characterize the relationship between sampling history and information valuation. Finally, the sample was drawn from university students at a single institution. Exploration strategies change across development (Giron et al., 2023; Harms et al., 2024; Zhuang et al., 2023), so the findings may not generalize to all age groups and settings. Replication in more diverse samples will be important for establishing the generalizability and precision of the present findings.

In sum, higher WMC was associated with more directed exploration, but not with random exploration or general choice consistency. This difference was accompanied by stronger discounting of previously sampled options (larger ω), which statistically accounted for the association between WMC and directed exploration in an exploratory mediation analysis. At the neural level, the frontocentral P2 on directed exploration trials was larger in the high-WMC group, suggesting that group differences are already present during early option evaluation. These findings point to the use of sampling history as a specific route through which cognitive capacity relates to exploration. Whether this relationship is causal, and how ω relates to early neural activity, remain to be tested in designs that manipulate capacity and combine neural and computational measures in one model.

## Data Availability Statement

The data supporting the findings of this study are available upon request from the corresponding author.

## Acknowledgments and Funding Information

This study was supported by the Guangdong Philosophy and Social Science Planning Project (Grant No. GD25YXL08) and the National Natural Science Foundation of China (Grant Nos. 32271139 and 32671451).

## Diversity in Citation Practices

Retrospective analyses of citation practices in cognitive neuroscience, including in this journal, have shown that papers from nonmale-identifying research groups are undercited relative to their proportions of authorship (Fulvio et al., 2021). We aimed to cite relevant work in a way that reflects the breadth of the field.

