## Supplementary Materials for "Working memory capacity is linked to directed exploration: behavioral, computational, and electrophysiological evidence"

### S1. Additional Behavioral Results under Reward Conditions

Additional analyses examined strategy use across low- and high-reward conditions under the unequal-information condition. A nonparametric analysis using the aligned rank transform (ART) was conducted to evaluate the effects of Reward Condition, Strategy, and Working Memory (WM) Group, together with their interactions.

**Table S1. Nonparametric ANOVA Results for Strategy Use across Reward Conditions under the Unequal Information Condition**

| Effect | F | df | Residual df | p |
|---|---|---|---|---|
| Reward Condition | 0.054 | 1 | 250 | 0.817 |
| Strategy | 123.87 | 2 | 250 | < 0.001 |
| WM Group | 0.083 | 1 | 50 | 0.774 |
| Condition × Strategy | 40.07 | 2 | 250 | < 0.001 |
| Condition × WM Group | 0.188 | 1 | 250 | 0.665 |
| Strategy × WM Group | 14.32 | 2 | 250 | < 0.001 |
| Condition × Strategy × WM Group | 0.098 | 2 | 250 | 0.906 |

**Note.** Reward Condition = low versus high reward; Strategy = exploitation, random exploration, and directed exploration. Results are based on Type III Wald F-tests using the Kenward–Roger approximation.

The analysis showed a significant main effect of Strategy and significant Condition × Strategy and Strategy × WM Group interactions. Reward Condition, WM Group, Condition × WM Group, and the three-way interaction were not significant.

**Table S2. Post-Hoc Comparisons of Strategy Use within Reward Condition under the Unequal Information Condition**

| Condition | Strategy | Mean (Low WM) | Mean (High WM) | t (df) | p | 95% CI of Difference |
|---|---|---|---|---|---|---|
| Low Reward | Exploitation | 0.441 | 0.323 | −1.83 (49.8) | 0.074 | [−0.248, 0.012] |
| Low Reward | Directed Exploration | 0.428 | 0.582 | 2.03 (49.7) | 0.048 | [0.001, 0.306] |
| Low Reward | Random Exploration | 0.108 | 0.085 | −0.71 (47.8) | 0.482 | [−0.089, 0.042] |
| High Reward | Exploitation | 0.695 | 0.559 | −2.50 (47.3) | 0.016 | [−0.245, −0.027] |
| High Reward | Directed Exploration | 0.146 | 0.331 | 2.94 (40.2) | 0.005 | [0.058, 0.312] |
| High Reward | Random Exploration | 0.128 | 0.097 | −1.01 (48.9) | 0.319 | [−0.092, 0.031] |

**Note.** Means represent the proportion of trials classified as each strategy. p values are two-tailed. Confidence intervals are for the between-group difference (High WM − Low WM).

**Figure S1. Strategy Use across Reward Conditions under the Unequal Information Condition**

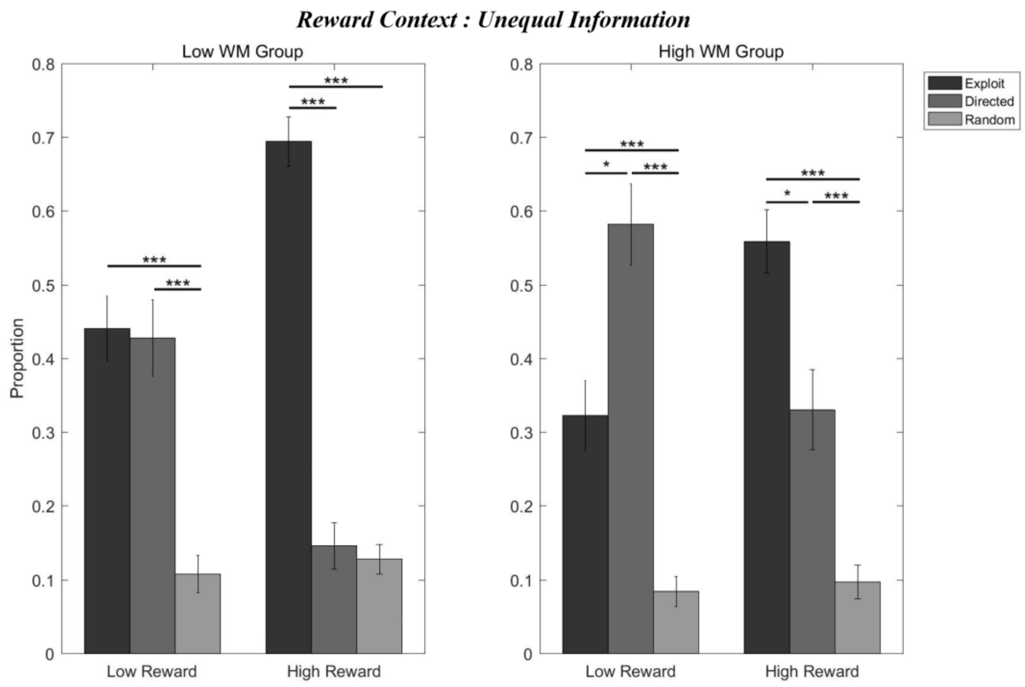

### S2. Additional ERP Analyses

An additional ERP analysis examined the exploration-versus-exploitation contrast at the Pz electrode across three analysis strategiess. Mauchly's test indicated a violation of sphericity for both the strategies main effect and the Group × Strategies interaction; therefore, Greenhouse–Geisser and Huynh–Feldt corrected results were examined.

**Table S3. Repeated-Measures ANOVA Results for the Additional ERP Analysis at Pz**

| Effect | df | F | p (GG) | p (HF) | ges |
|---|---|---|---|---|---|
| Group | 1, 50 | 1.070 | — | — | 0.014 |
| Strategies | 2, 100 | 9.464 | < 0.001 | < 0.001 | 0.057 |
| Group × Strategies | 2, 100 | 2.169 | 0.132 | 0.131 | 0.014 |

**Note.** Mauchly's W = 0.725 for both Strategies and Group × Strategies, ps < 0.001. Greenhouse–Geisser epsilon = 0.784 and Huynh–Feldt epsilon = 0.805 for both effects. ges = generalized eta squared. The omnibus Group × Strategies interaction was not significant.

The corrected analysis revealed a significant main effect of Strategies, whereas neither the Group main effect nor the Group × Strategies interaction was significant. Bonferroni-adjusted pairwise comparisons showed that exploitation differed from directed exploration (estimate = −2.98, $SE$ = 0.696, $t(102) = -4.288$, $p$ = 0.0001) and exploitation differed from random exploration (estimate = −1.70, $SE$ = 0.696, $t(102) = -2.436$, $p$ = 0.0498); the comparison between directed exploration and random exploration was not significant (estimate = 1.29, $SE$ = 0.696, $t(102) = 1.853$, $p$ = 0.2004).

**Table S4. Between-Group Comparisons at Each ERP Analysis Strategies**

| Strategies | Estimate (HighWMC − LowWMC) | SE | df | t | p |
|---|---|---|---|---|---|
| Exploitation | 0.00754 | 1.41 | 97.3 | 0.005 | 0.9957 |
| Directed exploration | 2.78985 | 1.41 | 97.3 | 1.986 | 0.0499 |
| Random exploration | 0.80008 | 1.41 | 97.3 | 0.569 | 0.5704 |

**Note.** Pairwise group comparisons were obtained from estimated marginal means. The original analysis used Bonferroni adjustment.

### S3. Computational Model Comparison and Parameter Recovery

The same model-selection rank order (gkRL > kRL > sRL) was observed within both working-memory groups, supporting the robustness of model selection. Parameter recovery analyses indicated adequate recovery for the remaining parameters, including ω, but poor identifiability for γ.

**Table S5. Between-Group Comparisons and Parameter Recovery Results for the gkRL Model**

| Parameter | Group | Group comparison | | | Parameter recovery | | | |
|---|---|---|---|---|---|---|---|---|
| | | **M** | **t** | **p** | **r** | **95% CI** | **RMSE** | **Quality** |
| ***α*** | High | 0.582 | −0.44 | 0.659 | 0.837 | [0.666, 0.925] | 0.084 | Good |
| | Low | 0.603 | | | 0.926 | [0.841, 0.967] | 0.071 | Excellent |
| ***β*** | High | 0.192 | 1.14 | 0.262 | 0.966 | [0.926, 0.985] | 0.026 | Excellent |
| | Low | 0.167 | | | 0.966 | [0.924, 0.985] | 0.016 | Excellent |
| ***ω*** | High | 8.22 | 2.90 | 0.007 | 0.988 | [0.974, 0.995] | 1.505 | Excellent |
| | Low | 2.51 | | | 0.904 | [0.795, 0.956] | 1.742 | Good |
| ***γ†*** | High | 0.035 | −10.12 | < 0.001 | 0.219 | [−0.184, 0.559] | 0.023 | Poor |
| | Low | 0.090 | | | −0.176 | [−0.528, 0.227] | 0.058 | Poor |

**Note.** M = posterior mean; t = independent-samples t-test statistic; p = two-tailed significance level. r = Pearson correlation between true and recovered parameter values; RMSE = root mean square error. Quality ratings: Excellent ($r \geq 0.90$), Good ($0.70 \leq r < 0.90$), Poor ($r < 0.50$). High-WMC and Low-WMC refer to the high and low working-memory-capacity groups, respectively. †γ showed a statistically significant between-group difference but was poorly identifiable; this difference should therefore be interpreted with caution.

### S4. Posterior Predictive Checks

Posterior predictive checks evaluated whether the gkRL model, fitted separately within each working-memory group, could reproduce the observed proportions of exploitation, directed exploration, and random exploration. Simulated and observed strategy proportions were moderately to strongly correlated across strategies (r = 0.561–0.789), with predictive precision indicated by RMSE values of 0.056–0.173.

Predictive performance was somewhat better in the High-WMC group, with stronger observed–predicted correlations and smaller RMSE values for random exploration. In the Low-WMC group, the model showed small discrepancies for exploitation and directed exploration, with mean observed–predicted differences of 0.075 and −0.066, respectively. Random exploration showed a smaller discrepancy (mean difference = −0.009). In the High-WMC group, mean differences were small across all three strategies (−0.024 to 0.023).

**Table S6. Posterior Predictive Checks for Strategy Proportions across Working Memory Groups**

| WM Group | Strategy | r | p (correlation) | RMSE |
|---|---|---|---|---|
| Low-WMC | Exploitation | 0.597 | 0.0013 | 0.156 |
| Low-WMC | Directed Exploration | 0.561 | 0.0029 | 0.173 |
| Low-WMC | Random Exploration | 0.575 | 0.0026 | 0.075 |
| High-WMC | Exploitation | 0.677 | < 0.001 | 0.166 |
| High-WMC | Directed Exploration | 0.789 | < 0.001 | 0.169 |
| High-WMC | Random Exploration | 0.697 | < 0.001 | 0.056 |

**Note.** *r* = Pearson correlation between observed and simulated strategy proportions; RMSE = root mean square error; Mean Difference = observed human proportion minus model-predicted proportion.

### S5. Full Mediation Model Parameters and Convergence Diagnostics

The full Bayesian mediation analysis consisted of four models. Path A estimated the association between working memory group (WMC group, low vs. high) and log-transformed ω. Path B estimated the association between log(ω) and strategy use, with exploitation as the reference category. Path C estimated the total association between WMC group and strategy use, and Path C′ estimated the direct association between WMC group and strategy use after controlling for log(ω). All Bayesian models were estimated using four chains with 4,000 iterations per chain, including 2,000 warmup iterations, yielding 8,000 post-warmup draws. All reported R-hat values were 1.00, indicating satisfactory convergence.

An earlier version of Path B was fitted using the raw ω parameter rather than its log-transformed value. This model was treated as a sensitivity analysis only and was not used as the formal mediation path. The final mediation model used log(ω) as the mediator to provide a more suitable representation of the parameter for the mediation analysis and to reduce the influence of the highly skewed raw ω scale. The raw-ω model is therefore not reported in detail here.

**Table S7. Full Bayesian Mediation Model Parameters and Convergence Diagnostics**

| Path / Model | Parameter | Estimate | SE | 95% CrI | R-hat | Bulk ESS | Tail ESS |
|---|---|---|---|---|---|---|---|
| Path A | Intercept | −0.60 | 0.35 | [−1.29, 0.09] | 1.00 | 6855 | 5039 |
| Path A | WMC group (High vs. Low) | 1.80 | 0.50 | [0.82, 2.77] | 1.00 | 6762 | 4961 |
| Path A | σ | 1.80 | 0.18 | [1.49, 2.20] | 1.00 | 7202 | 6123 |
| Path B | DE vs. Exploitation: log(ω) | 0.36 | 0.04 | [0.28, 0.45] | 1.00 | 3466 | 3925 |
| Path B | RE vs. Exploitation: log(ω) | −0.07 | 0.05 | [−0.16, 0.03] | 1.00 | 2568 | 4093 |
| Path C | DE vs. Exploitation: WMC group | 0.60 | 0.25 | [0.10, 1.10] | 1.00 | 1168 | 2130 |
| Path C | RE vs. Exploitation: WMC group | −0.17 | 0.20 | [−0.57, 0.21] | 1.00 | 1919 | 3642 |
| Path C′ | DE vs. Exploitation: WMC group | −0.02 | 0.18 | [−0.40, 0.33] | 1.00 | 2510 | 3755 |
| Path C′ | DE vs. Exploitation: log(ω) | 0.36 | 0.05 | [0.27, 0.46] | 1.00 | 2696 | 4143 |
| Path C′ | RE vs. Exploitation: WMC group | −0.06 | 0.22 | [−0.48, 0.37] | 1.00 | 2047 | 3940 |
| Path C′ | RE vs. Exploitation: log(ω) | −0.06 | 0.06 | [−0.17, 0.05] | 1.00 | 2133 | 3774 |

**Note.** DE = directed exploration; RE = random exploration; WMC group = working memory capacity group (low vs. high, high coded as the contrast level); CrI = credible interval; ESS = effective sample size. Intercepts, σ, and random-intercept standard deviations are reported as model parameters and convergence diagnostics rather than theoretical mediation paths. Random-intercept SDs were 0.49 [0.35, 0.66] for DE and 0.62 [0.47, 0.80] for RE in Path B; 0.85 [0.65, 1.09] for DE and 0.63 [0.49, 0.83] for RE in Path C; and 0.50 [0.36, 0.67] for DE and 0.62 [0.48, 0.81] for RE in Path C′. All corresponding R-hat values were 1.00.

As a frequentist cross-check of the Bayesian Path A analysis, a linear regression of log(ω) on WMC group (high vs. low) yielded a statistically significant positive association, $t = 3.69$, $p < .001$, $R^2 = .214$, adjusted $R^2 = .198$. This analysis was conducted as a robustness check and was not used as the primary mediation model.

The posterior distribution of the indirect effect was positive, with an estimate of 0.643 (95% CrI [0.284, 1.045]) and a posterior probability of 0.9999 that the indirect effect was greater than zero. For directed exploration, the corresponding odds ratio was 1.431 (95% CrI [1.319, 1.562]), with posterior probability 1.00 that the odds ratio exceeded 1.

**Table S8. Indirect Effect and Odds-Ratio Summary**

| Quantity | Estimate / Mean | Median | 95% CrI | Posterior Probability |
|---|---|---|---|---|
| Indirect effect | 0.643 | 0.635 | [0.284, 1.045] | P(indirect effect > 0) = 0.9999 |
| OR for DE | 1.431 | 1.428 | [1.319, 1.562] | P(OR > 1) = 1.00 |

**Note.** DE = directed exploration. The odds ratio was calculated from posterior draws for the directed-exploration coefficient.

Consistent with the primary mediation analysis, log(ω) and WMC were not associated with random exploration relative to exploitation.

### S6. Random Exploration Comparisons

To provide the complete set of mediation-model contrasts, the following results summarize associations with Random Exploration relative to Exploitation. None of the corresponding 95% credible intervals excluded zero.

**Table S9. Random Exploration Contrasts Relative to Exploitation**

| Path | Contrast | Estimate | 95% CrI | Interpretation |
|---|---|---|---|---|
| B | log(ω) → RE vs. Exploitation | −0.07 | [−0.16, 0.03] | CrI includes 0 |
| C | WMC → RE vs. Exploitation | −0.17 | [−0.57, 0.21] | CrI includes 0 |
| C′ | WMC → RE vs. Exploitation | −0.06 | [−0.48, 0.37] | CrI includes 0 |
| C′ | log(ω) → RE vs. Exploitation | −0.06 | [−0.17, 0.05] | CrI includes 0 |

**Note.** WMC = working memory capacity; RE = Random Exploration. Estimates are on the categorical-model log-odds scale relative to Exploitation.

### S7. Rescaling in Nested Categorical Models

In nonlinear models such as logistic or multinomial logistic regression, coefficients from nested models are not directly comparable in the same way as in linear regression, because the variance of the underlying latent variable is rescaled whenever a covariate is added (Mood, 2010; Karlson et al., 2012). This rescaling problem specifically concerns the comparison between the total effect (Path C) and the direct effect (Path C′), and raises the possibility that part of the observed attenuation from c to c′ reflects rescaling of the latent variable rather than genuine mediation. This concern applies less directly to the product-of-coefficients estimate of the indirect effect itself, which does not rely on comparing coefficients across nested models.

As a partial check, we compared the total effect to the sum of the direct and indirect effects, $c' + ab = -0.02 + 0.64 = 0.62$, which closely approximated the estimated total effect, $c = 0.60$. Because the linear decomposition $c = c' + ab$ is not guaranteed to hold exactly under a nonlinear link function, this close correspondence suggests that rescaling did not substantially distort the present pattern of results, although it does not rule out a modest contribution.

### S8. Correlations between Computational Parameters and P2 Amplitude

Additional exploratory analyses examined the associations between computational parameters and P2 amplitude across the overall sample and separately within the low- and high-WM groups. Pearson and Spearman correlation analyses were conducted to assess linear and monotonic associations, respectively. These analyses were performed to examine whether individual differences in computational parameters, particularly ω, were associated with variation in early neural activity during option evaluation.

**Table S10. Correlations between Computational Parameters and P2 Amplitude**

| Parameter | WM Group | *n* | Pearson *r* | *p* | Spearman ρ | *p* |
|---|---|---|---|---|---|---|
| α | Overall | 52 | 0.088 | 0.533 | −0.004 | 0.978 |
| α | High WM | 26 | −0.055 | 0.789 | −0.001 | 0.997 |
| α | Low WM | 26 | 0.276 | 0.172 | 0.161 | 0.432 |
| β | Overall | 52 | 0.071 | 0.616 | −0.034 | 0.810 |
| β | High WM | 26 | 0.073 | 0.721 | 0.105 | 0.611 |
| β | Low WM | 26 | −0.117 | 0.570 | −0.132 | 0.520 |
| γ | Overall | 52 | −0.430 | 0.001 | −0.507 | < 0.001 |
| γ | High WM | 26 | −0.049 | 0.813 | −0.169 | 0.410 |
| γ | Low WM | 26 | −0.109 | 0.595 | −0.209 | 0.306 |
| ω | Overall | 52 | 0.072 | 0.610 | 0.211 | 0.134 |
| ω | High WM | 26 | −0.238 | 0.241 | −0.210 | 0.304 |
| ω | Low WM | 26 | 0.080 | 0.698 | 0.178 | 0.385 |
| log(ω) | Overall | 52 | 0.101 | 0.474 | 0.211 | 0.134 |
| log(ω) | High WM | 26 | −0.286 | 0.156 | −0.210 | 0.304 |
| log(ω) | Low WM | 26 | 0.084 | 0.685 | 0.178 | 0.385 |

**Note.** Pearson correlations quantify linear associations between computational parameters and P2 amplitude, whereas Spearman correlations assess monotonic associations. Correlations were calculated across the overall sample and separately within the high- and low-WM groups. WM = working memory; P2 amplitude was measured at FCz during the early option-evaluation period.
